# The fog of genetics: Known unknowns and unknown unknowns in the genetics of complex traits and diseases

**DOI:** 10.1101/553685

**Authors:** João Pedro de Magalhães,, Jingwei Wang

## Abstract

Associating genetic variants with phenotypes is not only important to understand the underlying biology but also to identify potential drug targets for treating diseases. It is widely accepted that for most complex traits many associations remain to be discovered, the so-called “missing heritability.” Yet missing heritability can be estimated, it is a known unknown, and we argue is only a fraction of the unknowns in genetics. The majority of possible genetic variants in the genome space are either too rare to be detected or even entirely absent from populations, and therefore do not contribute to estimates of phenotypic or genetic variability. We call these unknown unknowns in genetics the “fog of genetics.” Using data from the 1000 Genomes Project we then show that larger genes with greater genetic diversity are more likely to be associated with human traits, demonstrating that genetic associations are biased towards particular types of genes and that the genetic information we are lacking about traits and diseases is potentially immense. Our results and model have multiple implications for how genetic variability is perceived to influence complex traits, provide insights on molecular mechanisms of disease and for drug discovery efforts based on genetic information.

## MAIN TEXT

Genetic variability contributes to nearly all human traits and diseases. The heritability of phenotypes, that is, the impact of genetic variation (versus environmental variation), can be small or large, and understanding the heritability of human traits is a major focus of modern genetics (Reich & Lander 2001; McClellan & King 2010; Timpson *et al*. 2018). Discovering genetic variants that are associated with disease risk and/or clinical outcomes is also extremely important. Genetic variants associated with diseases can inform about disease etiology, may have clinical applications and often become key therapeutic targets (Nelson *et al*. 2015; Visscher *et al*. 2017). Not surprisingly, large-scale genetic studies, including genome wide association studies (GWAS), are a major tool with >3,000 publications in the GWAS catalog (MacArthur *et al*. 2017) as of writing. However, genetic association studies can only detect associations for genes with existing functional genetic variants. Here, we propose that the finite genetic variability of the human species means we have a small genetic search space for association studies, which imposes constraints on our understanding of the genetic and molecular basis of human phenotypes. Indeed, we show that a greater genetic variability in a given gene correlates with a higher probability of identifying genetic associations in GWAS. We propose that there is a much greater unknown space of genotypes and phenotypes that remains undetected and even undetectable. Inspired by the term “fog of war”, referring to uncertainty in military operations, we call the uncertainty in genetics resulting from the limited natural variation of our species the “fog of genetics”.

As we try to understand the genetic basis of human traits and diseases, it is clear that we are missing extensive important information. It is well-established that for most phenotypes there is a significant element of missing heritability (Manolio *et al*. 2009; McClellan & King 2010; Zuk *et al*. 2012), that is, contributions from unknown genetic variants that explain variation in a phenotype like height, longevity or susceptibility to disease. Such missing heritability can be classified as the known unknowns of genetics. For a given phenotype we can estimate its heritability and, at least in theory but often in practice, estimate the impact of genetic variants. Therefore, even if for many phenotypes we are unaware of which and how many genes contribute to them, we can know how much of these genetic contributions we are missing, and indeed some use the term “hidden heritability” (Gibson 2010). However, there is also a much greater unknown in genetics: an ocean of unknown unknowns. On one hand, extremely rare variants can give rise to extreme phenotypes that are so rare they do not contribute to estimates of phenotypic variability. Some of these have been identified, for example in *PCSK9* affecting LDL cholesterol levels (Timpson *et al*. 2018), but how many are still unknown to us? That is unknowable at present, yet at least such variants exist in the human species.

If due to chance or selection a given allele becomes fixed in a species it stops being a genetic variant, no matter how big of a phenotypic impact it has compared to its now extinct variant(s). Even in the absence of genetic and phenotypic variability, the gene product in question will still play a major biological role in a given phenotype. Traits and diseases can be inherited even if they are not heritable. Besides, fixed variants may be the most important determinants of the phenotype of a species if they are adaptive and became fixed due to positive selection. Nonetheless, without a genetic association due to the absence of variants, important genes in phenotypes will go undetected. Because such genes are unknown and unknowable at present (but see below), they represent unknown unknowns in genetics. In other words, there is a huge space of potential genetic variation that we are unaware of because it does not presently exist in the human species. To use a military analogy, missing heritability is when we know the size and capacity of the enemy army, we just do not know its location. The unknown unknowns, the fog of genetics is when we do not even know the size or capacity of the enemy army.

It is certainly conceivable that a large number of phenotypically relevant genetic variants exist in humans yet they will be so rare as to go unnoticed. The human genome has roughly 3 billion base pairs, and each person has twice that amount of genetic material. But genetic variants have only been detected in a small fraction of the human genome sequence (Auton *et al*. 2015). For example, whole genome sequencing data from the 1000 Genomes Project revealed ~88 million variants (Auton *et al*. 2015). If one assumes a world population of 7 billion people, with 70 de novo mutations per individual (Kong *et al*. 2012), by chance mutations occurred in every single human nucleotide. Of course mutations that are lethal at embryonic stages will not exist in populations. More importantly, genetic changes affecting one or a small number of individuals, families or populations will go unnoticed in GWAS and even in estimates of heritability.

For genetic variants that have a high penetrance and strong, early onset phenotypes (e.g., developmental abnormalities or catastrophic phenotypes) scientists and clinicians may be able to detect them in individual patients. Cases of Mendelian diseases, which can be caused by de novo mutations, will be identified in many cases. Yet most human traits and diseases have a complex, non-Mendelian genetic architecture. As such, we argue that the fog of genetics is widespread across most human phenotypes, in particular complex age-related diseases which are now the major killers in modern societies. Even genetic association studies with very large numbers of individuals have a limited ability to detect associations in very rare alleles. In addition, rare functional variants in drug target genes have been shown to be geographically localized, making them difficult to catalog (Nelson *et al*. 2012). Similarly, recent rare variants in family lineages may play an important role in human disease (Lupski *et al*. 2011). On top of genetic variants that are too rare to be associated with phenotypes are the unknowable genetic variants that have been lost and do not exist in the human species anymore (Figure 1).

**Figure 1:**
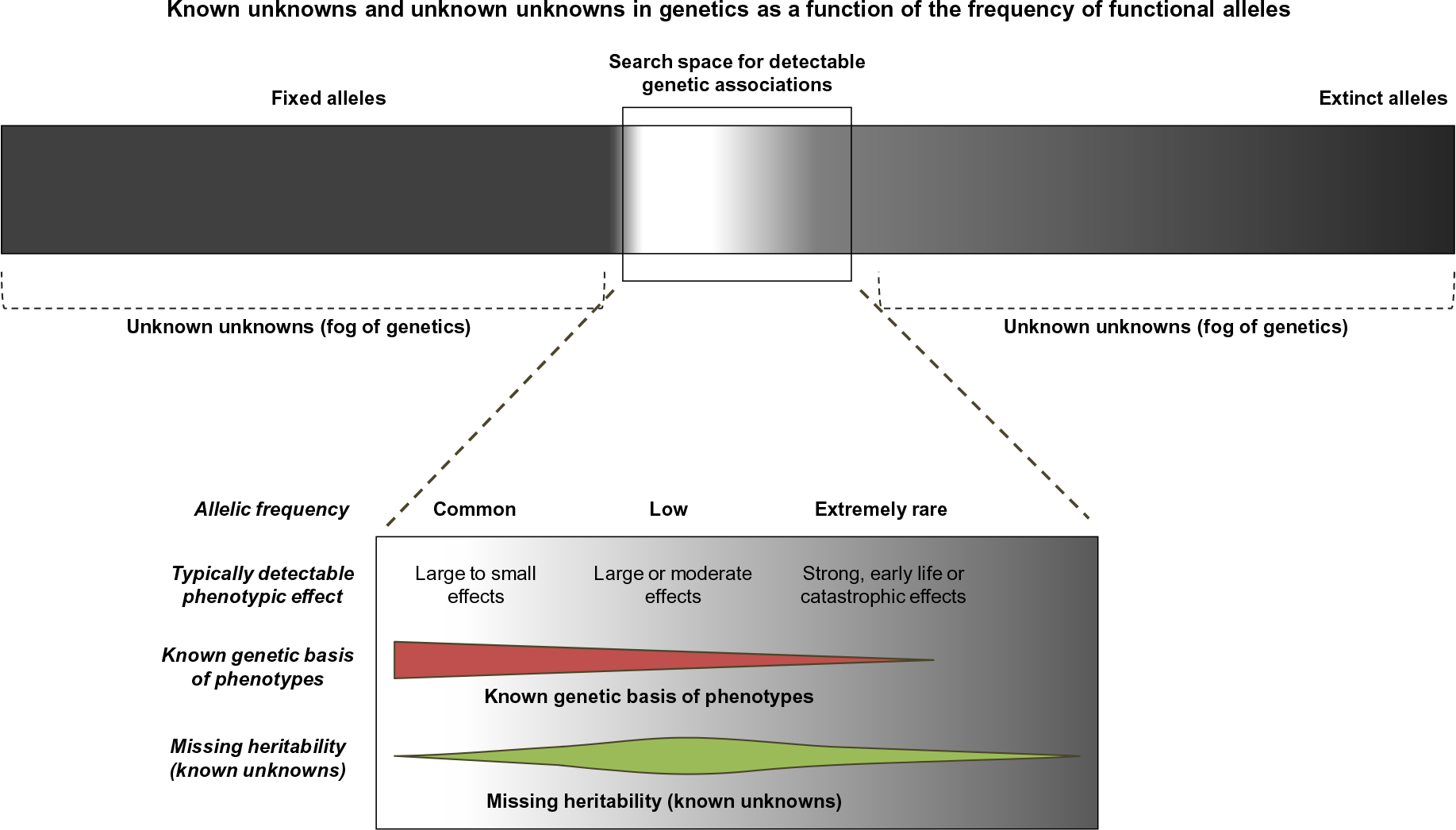
Detectability of phenotype-genotype non-Mendelian associations in relation to the frequency of genetic variants. Common variants, mostly ancestral alleles, can be associated with traits in GWAS. Alleles that have low frequencies in populations, however, will typically only be detected if they have strong phenotypic effects, like early onset diseases. At the extreme, genetic variants extinct in current populations (due to genetic drift or purifying selection) will not be detected. Likewise, alleles now fixed in populations will not be associated with phenotypes. In addition to the genetic variant frequency and strength of the phenotype, other factors not shown like penetrance and complexity of genetic architecture also affect detectability in genetic association studies.

Our framework predicts that the genes with more genetic variants will be more likely to be associated with phenotypes. To test this hypothesis, we estimated genetic diversity for each human gene based on the number of alternative alleles using data from the 1000 Genomes Project (Auton *et al*. 2015), normalized by gene length (see Supplementary Methods). We then counted the number of GWAS hits from the GWAS catalog (MacArthur *et al*. 2017) and analyzed the relationship between the number of GWAS hits and genetic diversity. As expected, genes with GWAS hits tend to be longer (Supplementary Table 1) and there is a correlation between gene size and number of GWAS hits (Kendall’s tau = 0.392 for protein-coding genes; tau = 0.260 for non-coding genes; p-value < 0.001 for both). In a way, this means that genetic association studies will be biased towards finding associations in larger genes, presumably because these have more genetic variants, even though larger genes will not *a priori* be the most important biologically. Larger genes also tend to have a slightly higher genetic diversity (Kendall’s tau = 0.057 for protein-coding genes; tau = 0.034 for non-coding genes; p-value <0.001 for both), we speculate because perhaps longer genes have a slightly lower chance of mutations being deleterious. Importantly, we found that genes with GWAS hits have a greater genetic diversity (Figure 2). Indeed, genes with a greater genetic diversity have more GWAS hits (Kendall’s tau = 0.106 for protein-coding genes; tau = 0.073 for non-coding genes; p-value < 0.001 for both), an effect that is still observed when accounting for the potentially confounding effects of gene length (p-value < 0.001). The differences observed are not huge (Figure 2; Supplementary Table 2), but given that most genetic variants are thought to be neutral (Kimura 1983), this is to be expected. Similar results were obtained using nucleotide diversity (π), another measure of DNA polymorphisms (Nei 1987), and in individual populations (Supplementary Results). Overall, these results demonstrate that genes with greater diversity have a higher probability of being associated with human phenotypes. Given that the power to detect a genetic association increases with the allelic frequency (Hong & Park 2012), our results are in line with theoretical expectations and crucially highlight that GWAS hits are biased towards particular types of genes.

**Figure 2:**
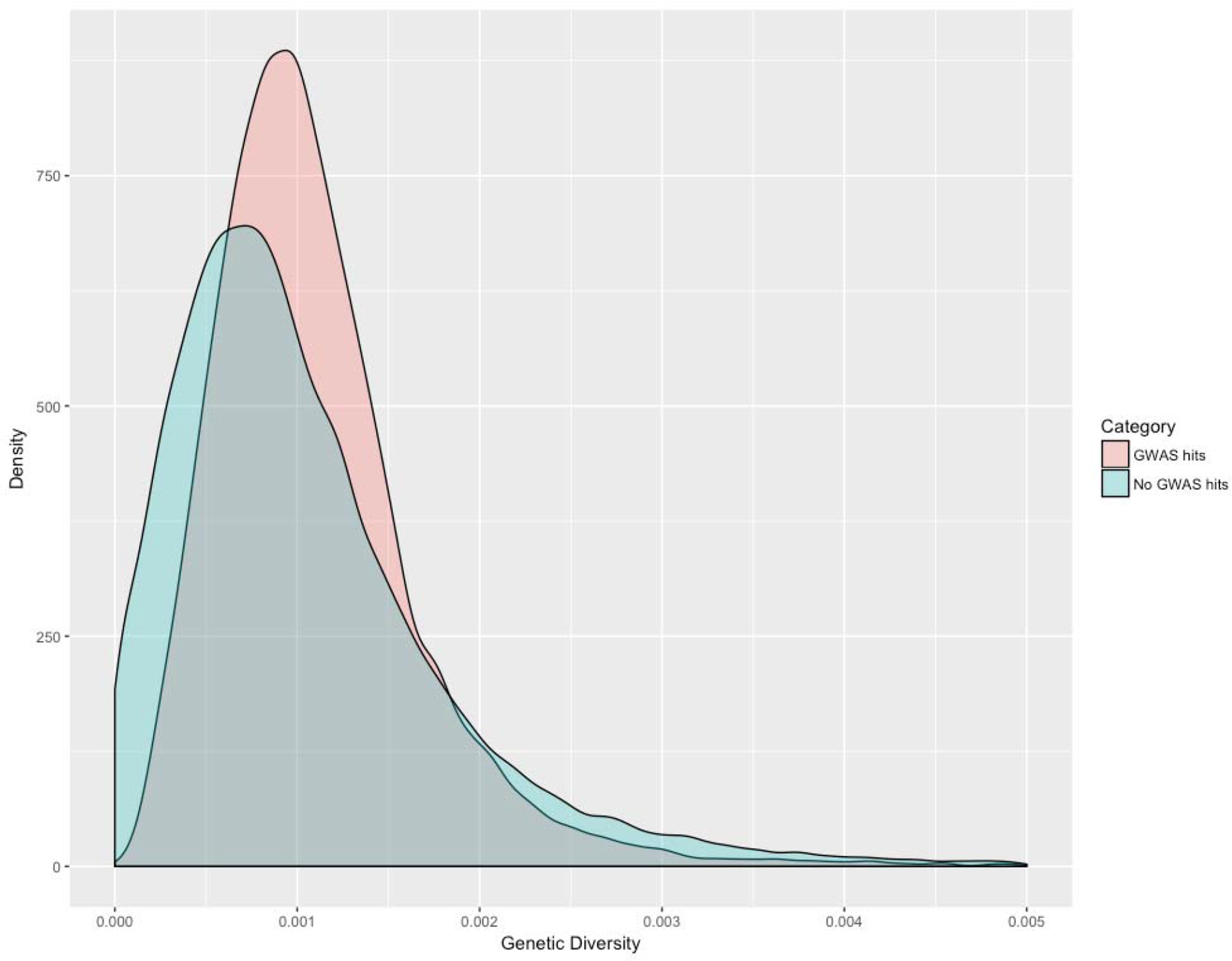
Density plot of genetic diversity of genes with hits in GWAS (pink; n =11,869) and genes without GWAS hits (green; n = 42,980). Only genetic diversity <0.005 were displayed. The difference between the genetic diversity of genes with hits in GWAS (median = 0.001009) and genes without GWAS hits (median = 0.000885) is highly statistically significant (p-value < 0.001; Mann-Whitney test). Please refer to *Supplementary Table 2* for further details of the genetic diversity of the two groups.

Rare genetic variants associated with several phenotypes have been detected (Timpson *et al*. 2018), and ongoing efforts aim to characterize rare disease-causing alleles (Saleheen *et al*. 2017). In addition, most genetic association studies employ European or Asian populations, and expanding genetic studies to other populations, in particular in Africa with its greater genetic diversity (which we also observed, see Supplementary Figure 1), will reveal novel functional genetic variants. Comparative genomics studies with other species, like primates (Muntane *et al*. 2018), can also uncover putative phenotypic effects in genetic variants that are fixed in humans. Besides, model systems and artificial forms of genetic diversity (mutagens, genetic screens, directed evolution, etc.) can help characterize the genetic landscape that is not naturally available, although by relying on model systems the relevance to human biology is not always straightforward. Nonetheless, these represent only a very small fraction of possible genotypes, and many of those unknown to us may be relevant for drug discovery efforts. Moreover, genetic variants conferring disease protection, particularly age-related diseases, will likely go unnoticed because not having a disease is not phenotypically exceptional. For instance, it has been traditionally difficult to associate genes with human longevity, in spite of huge advances in our understanding of the genetics of aging in model systems (de Magalhaes 2014). Lack of genetic associations with human longevity of aging-related genes could be due to a lack of functional variants in a complex phenotype. Indeed, most cancer driver genes initially were not known to be germline mutations predisposing to cancer (Vogelstein *et al*. 2013). Further experimental evidence for the fog of genetics comes from artificial selection experiments. Experimental evolution experiments in yeast show that, even though there is variation at the sequence level in different experiments, the same pathways seem to be selected for adaptation (Kryazhimskiy *et al*. 2014).

The fog of genetics predicts that the genetic basis of common complex traits and diseases entails only a fraction of the molecular players involved in the trait/disease; it thus follows that genes with an immense biological role in a given trait or disease may not necessarily be genetically associated with that trait/disease. Although it is obvious that genetic associations can only detect existing genetic variants and genetics only elucidates a fraction of the molecular events underlying a complex phenotype, these are overlooked issues, and we propose moving beyond natural genetic and phenotypic variation to understand complex phenotypes and diseases. There is clearly a huge number of unknown but possible genetic variants that could impact greatly on human traits and diseases. These unknowns will be smaller for high penetrance variants causing early onset, strong phenotypic effects and greater for late-onset complex diseases.

In conclusion, not only we are missing heritability of complex traits and diseases (the known unknowns of genetics), but we are unaware of a huge number of possible variants that either do not exist or are so rare in the human species that cannot be current associated with phenotypes (the unknown unknowns of genetics). In other words, missing heritability is only one component (and a small one, we argue) of what we do not know in genetics. We call the much we cannot learn from the natural variation of our species the fog of genetics. The implications of the fog of genetics are multiple and far-reaching: our molecular understanding of most diseases and traits is based on a very limited human genetic landscape. In a way, genetic variation is a species-specific phenomenon, our genetic diversity is a snapshot of our species’ long evolutionary journey. In fact, the genetic diversity of our species is lower than for other primates, like our closest-living relatives chimpanzees (Prado-Martinez *et al*. 2013). To put it another way, what we know of gene function in health and disease is based on the limited genetic and phenotypic natural variation of the human species; genetic associations are also biased towards the longer genes with the most variants. Indeed, the molecular components of phenotypes often do not carry genetic associations (Menche *et al*. 2015), and others have proposed that we must move beyond genes associated with diseases to understand disease etiology (Boyle *et al*. 2017). Even though unknown genetic variants that do not exist are not responsible for phenotypic variation (or clinical cases) these are still important because they can serve as target for therapeutics and can be crucial molecular players in biological processes and diseases. Lastly, current drug discovery efforts based on genetic variants (Nelson *et al*. 2015) employ a narrow and limited information, and expanding these by studying more diverse populations, artificial forms of generating genetic diversity and by improved phenotyping offers great promise for biomedical research.

## Acknowledgements

We are grateful to Saara Marttila, Yang Li, Zhengdong Zhang, Alessandro Cellerino and Arcadi Navarro for critical comments on previous versions of this manuscript and to Sònia Casillas Viladerrams for assistance obtaining nucleotide diversity data. Work in our lab is supported by the Wellcome Trust (104978/Z/14/Z and 208375/Z/17/Z), the Leverhulme Trust (RPG-2016-015), LongeCity, the Methuselah Foundation and the Biotechnology and Biological Sciences Research Council (BB/R014949/1).

