## Supplementary material for "The fog of genetics: Known unknowns and unknown unknowns in the genetics of complex traits and diseases"

**Contents**

*In this file*

**Supplementary Methods**
**Supplementary Results**
**Supplementary Table 1.** Gene length in GWAS-hit genes (GHGs) vs non-GWAS-hit genes (non-GHG).
**Supplementary Table 2.** GD in GWAS-hit genes (GHGs) vs non-GWAS-hit genes (non-GHG).
**Supplementary Figure 1.** Comparison of GD between five major populations

*In accompanying Excel file*

**Dataset S1.** Full list of genes with GD and GWAS hits

**Supplementary Methods:**

*Calculation of genetic diversity*

Variant calls were retrieved from the final phase (phase 3) of the 1000 Genomes Project (http://ftp.1000genomes.ebi.ac.uk/vol1/ftp/release/20130502/) (Auton *et al.* 2015). Genetic Diversity (GD), which was calculated from the 1000 Genomes Project global population data, refers to the nucleotide diversity between genes. The normalised (by length) minor allele frequency (MAF) of SNPs in each gene was used to represent GD (see below). Because GRCh37 is the reference genome used by the 1000 Genomes Project phase 3 variants, GRCh37 was used as the reference genome for genetic variants mapping. We calculated GD for genic regions including introns, given that most genetic associations are found in non-coding regions and intronic regions specifically (Freedman *et al.* 2011). As the regions in upstream and downstream often contain regulatory elements which could affect functions of the gene, the flanking regions 1000bp upstream and downstream of each gene, which tends to be under strong LD, were also included when mapping variants to genes and calculating GD . The start and stop positions of genic and flanking regions were retrieved from GRCh37 using Ensembl BioMart. Then, the genetic variants from the 1000 Genomes Project were mapped to genes containing genic and flanking regions. The calculation of GD was performed as follows:

1. Consider a cohort consisting of $n$ individuals, for a given SNP at position $i$, the total number of minor alleles (representing nucleotide changes) at position $i$ in the population is:

$$n \times MAF_i$$

2. If there are $m$ SNPs located in the genic region and two flank regions, the total number of minor alleles within the genic and flank regions can be calculated by:

$$\sum_{i=0}^{m} n \times MAF$$

3. As $n$ is a constant, the above formula can be simplified as:

$$\sum_{i=0}^{m} MAF$$

4. Then the normalised minor alleles in every 1000bp DNA is:

$$GD = \frac{1000 \times \sum_{i=0}^{m} MAF}{length_{genic} + length_{upstream} + length_{downstream}}$$

In total, 54,849 genes were obtained from the GRCh37 assembly, including protein coding gene, lincRNA, anatisense, misc_RNA, snRNA, pseudogene. See Dataset S1 for the full dataset, including GD values.

*Nucleotide diversity (π) data*

Data on nucleotide diversity (π) per gene was obtained for the whole genic region of protein-coding genes from PopHuman, a genomics browser based on the 1000 Genomes Project (Casillas *et al.* 2018). To focus on individual populations, π was obtained from three populations: Utah residents with Northern and Western European ancestry (CEU), Han Chinese in Beijing, China (CHB) and Yoruba in Ibadan, Nigeria (YRI).

*Counting the number of traits reported in the GWAS-Catalog per gene*

Data files were downloaded from the GWAS-Catalog website on 31/01/2018 (MacArthur *et al.* 2017). Then, in each entry, the reported SNPs were mapped to genes (with 1000bp in each upstream and downstream flanks, as for the GD calculations) in the GRCh37 assembly using rsIDs. SNPs mapping to two or more adjacent genes are counted separately. Through this method, connections between studied traits and genes (where reported SNPs located in) were recovered. By counting the number of unique *Mapped Traits* that are associated with any given gene, the corresponding number of associated traits of a gene was obtained.

*Statistical analyses*

Non-parametric tests were employed for the statistical analyses. The Mann–Whitney U-test was used to compare different groups of genes (e.g., genes with GWAS hits versus genes without GWAS hits). Correlation analyses were performed using Kendall's rank correlation (similar results were obtained using Spearman's rank-order correlation). Statistical analyses were performed using SPSS version 22 (IBM), R version 3.4.2 (R Core Team 2017) and RStudio version 1.1.383 (RStudio Team 2015).

**Supplementary Results:**

After merging π to the number of traits associated with each gene, we obtained a dataset with 21,020 genes. The analyses for GD reported in the main text were repeated using π from CEU, CHB and YRI with very similar results. For all three populations, genes with a greater π have more GWAS hits (Kendall's tau for CEU = 0.127; tau for CHB = 0.114; tau for YRI = 0.087; p-value < 0.001 for all three); the correlation between π and number of GWAS hits in a gene is still statistically significant after accounting for the effects of gene length (p-value < 0.001 for all three populations). Besides, genes with GWAS hits (GHGs) have a greater π than genes without GWAS hits (non-GHG): GHGs = 0.0006781 vs non-GHG = 0.0005552 for CEU; GHGs = 0.0006222 vs non-GHG = 0.0005142 for CHB; GHGs = 0.00088535 vs non-GHG = 0.0007907 for YRI (using a Mann-Whitney test, differences are highly statistically significant at p-value < 0.001 for all populations).

***Supplementary Table 1:*** *Gene length in GWAS-hit genes (GHGs) vs non-GWAS-hit genes (non-GHG).*

| Gene class | n | Min. | Median | Mean | Max. |
|---|---|---|---|---|---|
| Genome-Wide | 54849 | 2008 | 4913 | 32270 | 2306638 |
| GHGs | 11869 | 2047 | 47011 | 100576 | 2306638 |
| non-GHGs | 42980 | 2008 | 3291 | 13408 | 1231306 |
| Protein-coding GHGs | 8196 | 2186 | 59864 | 118152 | 2306638 |
| Protein-coding non-GHGs | 11234 | 2059 | 16368 | 30762 | 1231306 |
| Non-coding GHGs | 3673 | 2047 | 19777 | 61357 | 1538213 |
| Non-coding non-GHGs | 31759 | 2008 | 2688 | 7266 | 456257 |

***Supplementary Table 2.*** *GD in GWAS-hit genes (GHGs) vs non-GWAS-hit genes (non-GHG).*

| Gene class | n | Min. | Median | Mean | Max. |
|---|---|---|---|---|---|
| Genome-Wide | 54849 | 0 | 0.000923 | 0.001098 | 0.033518 |
| All GHGs | 11869 | 0.000024 | 0.001009 | 0.001158 | 0.033518 |
| All non-GHGs | 42980 | 0 | 0.000885 | 0.001075 | 0.019021 |
| Protein-coding GHGs | 8196 | 0.000061 | 0.000976 | 0.001073 | 0.031054 |
| Protein-coding non-GHGs | 11234 | 0 | 0.000846 | 0.000972 | 0.013428 |
| Non-coding GHGs | 3673 | 0.000024 | 0.001087 | 0.001347 | 0.033518 |
| Non-coding non-GHGs | 31759 | 0 | 0.000903 | 0.001112 | 0.019021 |

***Supplementary Figure 1.*** *Comparison of GD between five major populations: AFR, African; AMR, Ad Mixed American; EAS, East Asian; EUR, European; SAS, South Asian.*

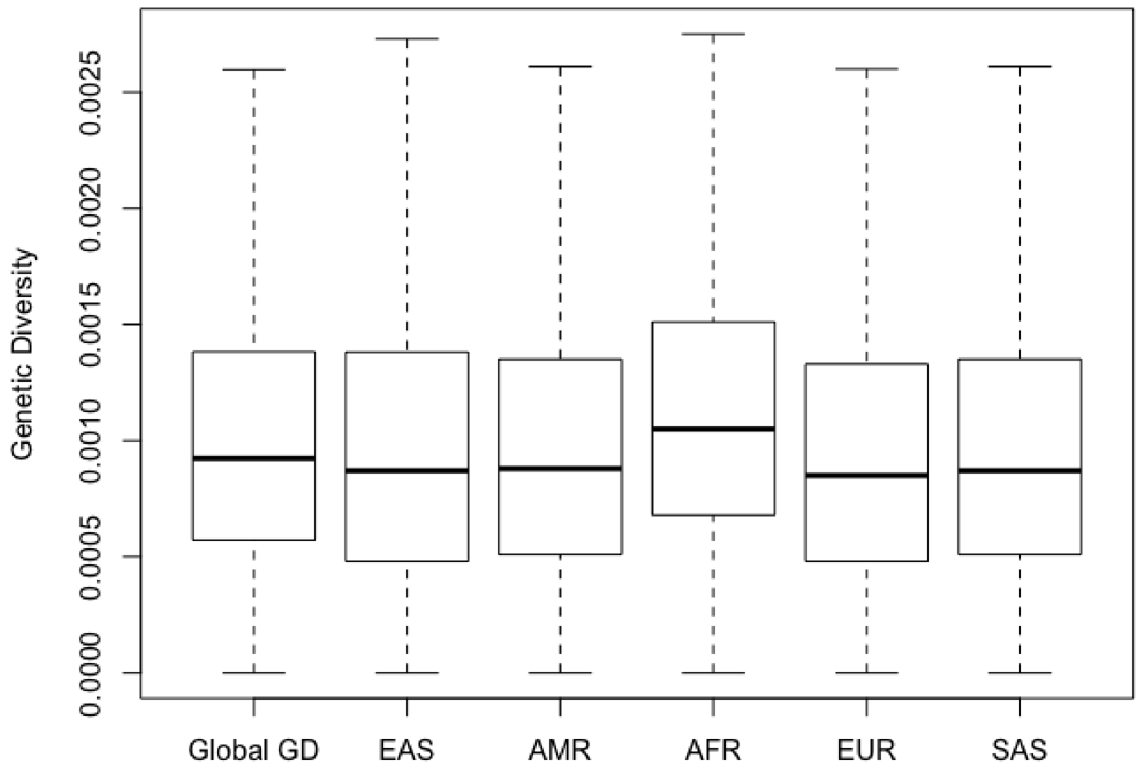
